# Potential for Biodiversity in Shared Residential Yards

**DOI:** 10.1101/2024.11.11.622952

**Authors:** Havi Livne, Michal Gruntman, Efrat Blumenfeld Lieberthal

**Affiliations:** Tel Aviv University

## Abstract

In urban environments, attention has traditionally been paid to public open spaces. However, private open spaces constitute a significant percentage of the urban area and can thus play a significant role in enhancing biological diversity and facilitating direct interactions between humans and nature within urban settings. According to the 2018 European Union report, 46% of homeowners across Europe live in apartment buildings and yet, most studies that examine the effect of private open spaces on biodiversity focus primarily on yards of single-family housing. The objective of this study is to examine the biodiversity potential offered by shared residential yards focusing exclusively on wild plant species, with the aim of offering planning guidelines that address both the management practices of these lot spaces and their spatial arrangement and qualities. The study was conducted in the city of Givatayim, Israel, in shared residential yards. The basic units (lots and buildings on them) in the city were delineated using a novel approach through a separate characterization of lot sizes and building types, as well as their combination. This process resulted in typologies that cover all residential fabric in the city, allowing systematic sampling. In the spring of 2022, plant surveys were conducted in 56 randomly chosen yards. In these lots, potential open area was measured, and the level of non-maintenance (neglection) was also rated. 74 species of wild plants were found, representing a quarter of all wild plant species in the city. Our results show the combined positive effect of available open space and levels of non-maintenance in these yards on the richness and diversity of wild species. This study highlights the substantial area occupied by residential yards in the city and its role in maintaining biodiversity in urban environments. This study indicates the importance of considering these aspects in architectural and management policies in cities.

## Introduction

Approximately 50% of the surface of the Earth is influenced by human activities. Urbanization, covering approximately 4% of the Earth’s surface, continues to be a predominant factor affecting natural ecosystems and stands as the most rapidly expanding form of land use (Loram et al., 2008). The interaction between humans and nature within urban contexts forms a reciprocal relationship, which is of primary focus within the domain of urban ecology. This discipline seeks to merge ecological science with other scientific and planning fields to enhance human living conditions while ensuring urban development remains compatible with ecological systems over the long term (Niemelä, 1999).

Urban environments generate extreme environmental conditions, challenging ecological theories originally derived from studies of more natural or less disturbed settings (Niemelä, 1999). The wide range of human activities leads to the degradation and fragmentation of habitats (Davis, 1978; Niemela, 1999; Clergeau et al., 2006; Gupta, 2002; Savard et al., 2000), with urbanization significantly contributing to the loss of existing and potential habitats (Mcdonald et al., 2008) and fostering environments where a limited set of species associated with humans flourish at the detriment of wild species (Hobbs et al., 2006).

Urban green infrastructure plays a critical role in mitigating urbanization’s adverse impacts on biodiversity by offering habitats more akin to natural conditions compared to “gray” urban or industrial areas (Filazzola et al., 2019). Adequate vegetated spaces within cities are therefore necessary to preserve diversity, underscoring urban planning’s role in transforming cities into sanctuaries for species threatened by agricultural practices.

Research indicates that the spatial arrangement and size of green spaces within urban green infrastructure significantly influence the city’s biological diversity. Dense urban development allows for the preservation of large, albeit isolated, green spaces (land-sparing), whereas dispersed development results in numerous fragmented small green spaces throughout the urban area (land-sharing) (Lin & Fuller, 2013). While some scholars advocate for land-sparing as the superior approach for biodiversity conservation (Soga et al., 2014), others argue that urban environments, characterized by habitat patches, offer diversity in smaller and more isolated areas, especially towards city centers (Mcdonald et al., 2008). Such spaces form permeable networks enhancing the urban landscape’s permeability (Shanahan et al., 2011), enabling human interaction with accessible and proximate nature (Miller & Hobbs, 2002), and delivering ecosystem services that improve residents’ well-being (Fuller et al., 2007).

Furthermore, recent studies identify private yards as potential habitats for wild native species (Berthon et al., 2021; Grade et al., 2021; Jimenez et al., 2022; Larson et al., 2020a), advocating for an integrated approach as the optimal strategy for preserving and enhancing biodiversity (Grade et al., 2021; Kremen, 2015; Valente et al., 2022), thereby emphasizing the significance of habitat size. For example, a meta-analysis on urban biodiversity in 2015, examining a wide range of taxonomic groups across 75 cities globally, revealed that habitat size and connectivity via corridors exert the most significant influence on biodiversity, enhanced by vegetation structure (Beninde et al., 2015). This study underscored that while biodiversity-friendly management has beneficial effects, expanding habitat areas and establishing corridor networks are paramount for sustaining high biodiversity levels in urban settings (Beninde et al., 2015).

Contrary to the conventional minimum habitat size standards for conservation, which suggest areas ranging from hundreds to thousands of dunams, Riva et al. (2023) presented evidence that even small habitats, from one to ten dunams, substantially contribute to the biodiversity of numerous species and ecosystems (Riva & Fahrig, 2023). This challenges the prevailing preference for larger over smaller areas, citing studies that demonstrate higher diversity within a collection of smaller spaces compared to a singular large area (Deane & He, 2018; Fahrig & Storch, 2020; Hammill et al., 2020), advocating for a paradigm shift in biodiversity conservation within human-dominated landscapes.

The concept of complementary ecological land uses, introduced by Colding (2007), highlights the synergistic potential of combined habitats to support biodiversity. This approach dissolves the traditional public-private area distinction, urging planners to view all spaces as a unified system (Colding, 2007). Residential yards, constituting a significant portion of urban land area, can enhance spatial connectivity and provide essential green spaces within cities. For instance, in Great Britain, yards account for an average of 23% of urban areas (Loram et al., 2008), and in Dunedin, New Zealand, up to 36% (Mathieu et al., 2007). Studies indicate these residential areas often contain the majority of urban greenery and vegetation (Ossola et al., 2019a; Ripplinger et al., 2017). However, despite their potential, conservation efforts frequently overlook residential yards in favor of nature reserves and parks, limiting the scope for biodiversity preservation in residential contexts (Lerman et al., 2023). Furthermore, interactions between yards and other green spaces have been documented, such as a study showing that private residential yards can boost bird species richness in adjacent public parks (Chamberlain et al., 2007). Yet, research on urban biodiversity often focuses on public green spaces rather than private yards (Baldock et al., 2019; Botzat et al., 2016), indicating a gap in the understanding and utilization of residential yards in urban biodiversity strategies.

The research that deals with residential yards in the last decade mostly focuses on yards owned by a single landowner rather than shared ones. Additionally, the research deals mainly with the characters and the actions of the landowners, from a perspective of management and maintenance, and the extent of their impact on biodiversity in the yard. As a result, there is a significant knowledge gap regarding the spatial characteristics of the yard itself that enhance biodiversity in residential yards, especially regarding the shared ones (Beumer & Martens, 2015; Blanchette et al., 2021; Gerner & Sargent, 2022a; Larson et al., 2022; Smallwood & Wood, 2023a).

In a pioneering study by Hoelzer et al. (2006) on private residences in Sheffield, UK, systematic relationships were identified between the size of individual housing lots and their yard extents, along with an exploration of the influence of house age and architectural style on yard characteristics. The findings indicated that larger yards tend to host a greater diversity of ground cover plants and trees (Hoelzer et al., 2006). Since plants form the basis of food for a wide network of consumers, yards containing higher proportions and abundance of wild plants support diverse and richer communities, contributing to biological diversity (Berthon et al., 2021; Coetzee et al., 2018; Lerman & Warren, 2011).

The vegetation and soil in open urban areas are crucial for supporting biodiversity, yet these elements are highly susceptible to alteration through human maintenance activities. Maintenance levels, including practices like mowing, pruning, and fertilizing, can have a negative impact on the diversity of birds, invertebrates, and insects in urban green spaces (Goddard et al., 2017; Kevin, 2008). For example, during the 2008 recession in the USA, a change in vegetation composition and an increase in annual wild species were observed in many yards as a result of halted irrigation and fertilization practices (Ripplinger et al., 2017). However, most studies that examine plants in residential yards, do not distinguish between wild plants and cultivars deliberately planted by landowners (Blanchette et al., 2021; Larsen & Harlan, 2006; Loram et al., 2008; Smallwood & Wood, 2023b; Wheeler et al., 2022b).

The nature of ownership of the yard and their maintenance practices have been shown to be crucial for biodiversity (Aronson et al., 2017; Colding, 2007; Shwartz et al., 2013). The shared property, described as an intermediate space between private and public, and therefor often lacks clear responsibility for maintenance (Rabinowitz, 2008). The situation is different in luxury towers with partially privatized common areas, whose residents treat these spaces as economic assets, integral to the property’s value (Azian et al., 2020; Wood et al., 2009). The levels of yard maintenance in shared ownership have changed over the years, as global wealth levels have increased and the property has been perceived as having economic value, the level of maintenance has also increased. According to Lerman et al. (2023), management decisions should ideally be taken at the neighborhood or district level in order to support a mosaic of habitats within the urban landscape. The reason for this recommendation is the numerous factors influencing maintenance methods (economic considerations, ownership, inclination towards nature conservation, aesthetics, and more), which create significant diversity in the field (Lerman et al., 2023).

Another factor that was shown to affect plant species richness in residential courtyards is their proximity to natural sites, which can increase the migration probabilities of propagules from the regional species pool (Muratet et al., 2007; Belaire et al., 2014). For example, in a study by Muratet et al. (2007) in Paris, neighboring large sites were found to be more floristically similar than distant sites, suggesting that the migration of species between large sites may significantly impact local plant composition. In contrast, small sites, which have a lower probability of receiving propagules by chance and are often isolated by built-up or impermeable areas, are less influenced by the composition of the surrounding sites.

This study examined the factors influencing biological diversity in **shared** residential yards. The choice to focus on shared residential yards stems primarily from the perception that the vast majority of research concentrates on private yards, and there is a significant knowledge gap regarding the potential for biodiversity inherent in shared yards, especially considering that they occupy a significant portion of the land surface for residential purposes, in many regions around the world. In 2019, 46% of the EU population lived in flats, while just over a third (35%) of the population lived in detached houses and almost one fifth (19%) lived in semi-detached or terraced houses (*House or Flat: Where Do You Live?*, 2021). Flats were the most common residence type in 14 Member States, notably in Latvia (where 66% of people lived in flats), Spain (65%) and Estonia (61%).

Secondly, the issue of non-maintenance, identified as a crucial element of this analysis, generates greater interest in spaces managed collectively by multiple individuals rather than those under individual stewardship. Moreover, in comparison to private yards, <u>shared</u> courtyards are characterized by a much wider variety of potential space types, as well as a significantly wider range of maintenance levels.

Here, we examined the effects of the size of shared residential yards, their levels of non-maintenance and distance to public green areas on the diversity of wild plants in a densely populated city. Unlike previous studies, we exclusively focused on wild plant species that have naturally colonized these yards rather than deliberately planted, to determine the parameters that influence the presence of natural biological diversity in these yards.

To represent all spatial configurations and fundamental units in the city, we characterized all lot and building sizes and their integration, thereby generating the architectural typologies of common buildings and their shared open spaces within the city. This methodology allowed us to seek connections between typology structural and functional characteristics and biodiversity.

## Methods

### Characteristics of the study area

The research focuses on the urban fabric of the city of Givatayim, Israel, which encompasses both historical and contemporary urban structures. Givatayim, ranking as Israel’s second most densely populated city, has a population density of 19,755 inhabitants per square kilometer (2023 municipality report). Findings from an urban nature survey (Mendelson, 2016) indicate that the majority of the city is situated on kurkar soil (a limestone that is uniquely found along the coastline), with sections in the southeast and north comprising clay soils, which are rare soil types in natural habitats and home to endemic plant and animal species. Givatayim is found in a Mediterranean climate characterized by moderate temperatures and high humidity. The climate is significantly influenced by the city’s closeness to the sea, which moderates both the diurnal and seasonal temperature variations. The region experiences hot, extended, dry summers and cool, mild, brief winters, with an average annual rainfall between 500-600 mm (*Israel Meteorological Service*, n.d.).

### Typology Characterization of Buildings and yards

In the urban context of Givatayim, non-public outdoor spaces are categorized into privately owned areas (such as balconies, gardens, and private rooftops) and those collectively owned by residents of a building or a complex (such as shared rooftops and yards). To create a survey representative of all spatial configurations and fundamental units (lots and the buildings upon them) in the city, we characterized all lot and building sizes and their integration, thereby generating typologies within the city. Traditionally, typo-morphological analysis is based on visual analysis of cartographic representations of streets, lots and/or buildings (Chen, 2012; Zhang, 2014). Urban typologies can focus on separate components of urban form, or combine several components and scales (Fusco & Araldi, 2017; Berghauser Pont et al., 2019).

All residential lots in the city (totaling 2,744) were measured, utilizing manual measurements of lot areas via AutoCAD software, based on GIS (Geographical Information Systems) data representing the original lots as provided by the Givataim municipality. These lots were then divided into four subcategories based on their sizes: 150-500, 500-800, 800-1500, and over 1500 m^2^ (Figure 1A). This categorization reflects the historical development patterns of Givatayim and aligns with patterns identified in the literature as characteristic of the city’s various zones.

**Figure 1:**
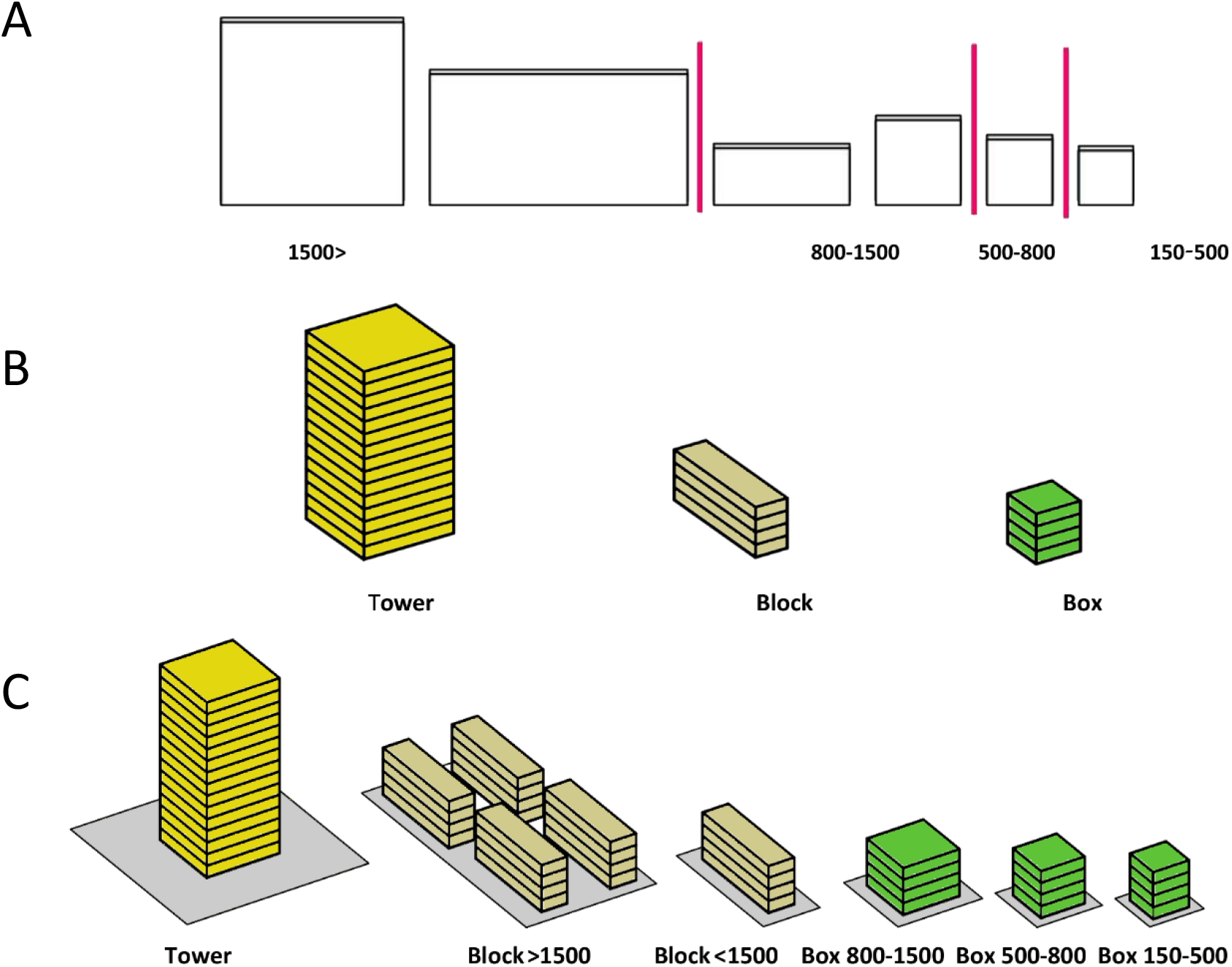
Typology categorization, including (A) segmentation of lots into four distinct size categories; (B) classification of building types into three distinct categories; and (C) synthesis of typologies as the combination of lot sizes and building types, which covers all shared residential areas.

The categorization of buildings within the city was conducted in a manner akin to that of lot types, drawing upon a synthesis of literature review and field observations. The categorization into building typologies aligns with conventions in scholarly research, grounded in analogous inquiries in the field (Hoelzer et al., 2006; Loram et al., 2008; Murdoch & Al-Habashna, 2024; Tooke et al., 2011). Given that the focal points of this investigation are the built and open cover within the lot and the interaction of the building with the ground level, buildings were classified into three primary archetypes: box buildings, which encompass all square structures up to 12 stories; elongated block buildings featuring 2-4 entry points; and towers of 12 stories and higher (Figure 1B). Consequently, six typologies emerged as combinations of building type and lot size category (Figure 1C), based on which all residential lots in the city were systematically mapped using Simplex 3D and GISNET V5 (Figure 2). The six typologies used in the research were proven to be effective for systematic mapping, as they covered all shared residential areas in the city, offering a reproducible approach for future studies.

**Figure 2:**
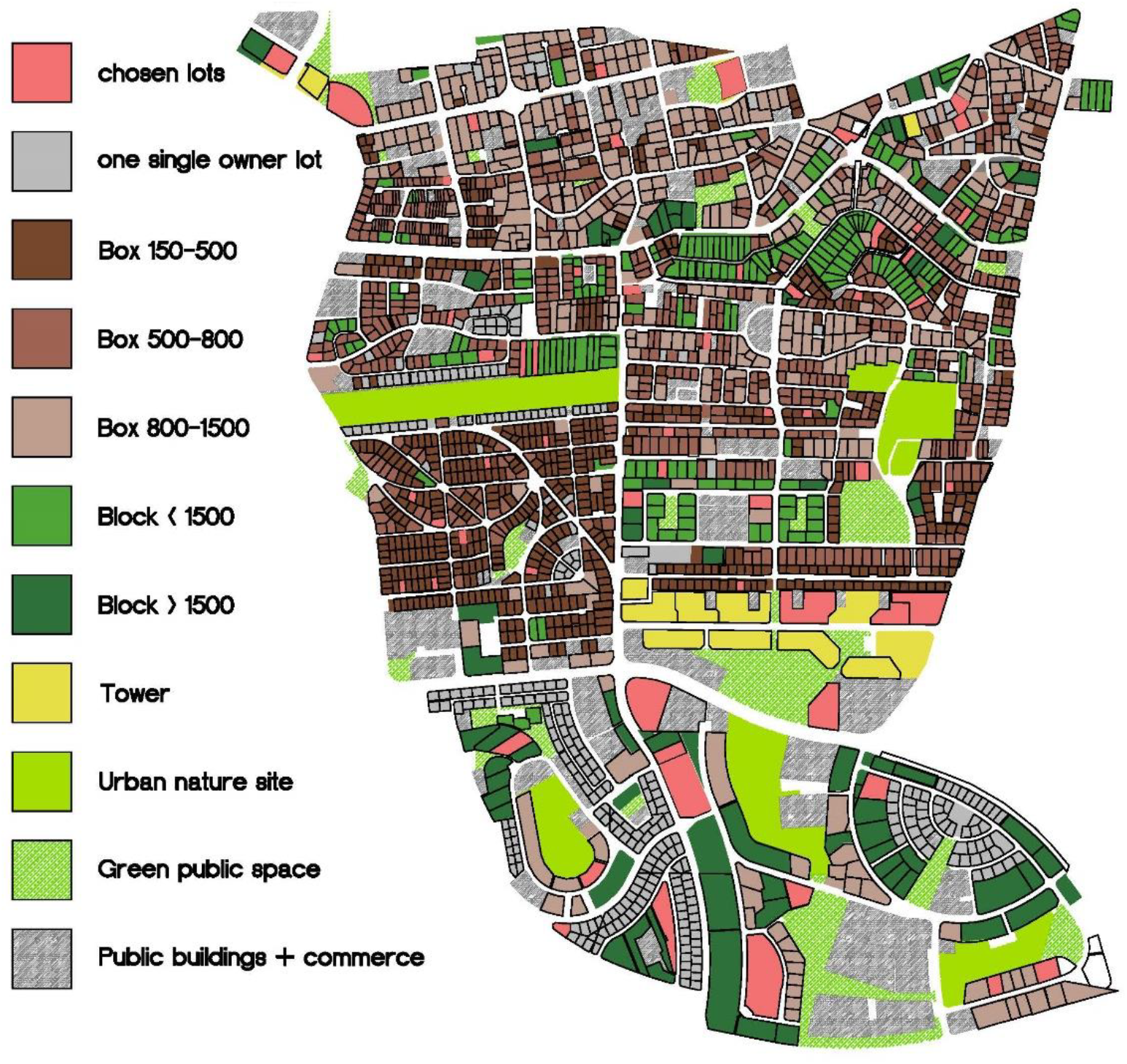
Mapping of all 2,477 residential lots within the city of Givatayim, categorized according to the six distinct typologies.

### Yards Selection and Characterization

Following mapping and classification of all residential lots within the city (Figure 2), 10 lots (yards) were randomly selected for each of the six typologies (60 lots in total) through a computerized lottery process, which utilized a code that specifically targeted the lot polygon. This code was developed using Grasshopper (an algorithmic modelling plugin for Rhinoceros® modelling software, version 6).

For each of the 60 selected lots, measurements were taken for three spatial parameters, including total lot area size, open area size (lot area uncovered by the building), and potential open area size (lot area uncovered by the building and impervious surfaces such as paths or parking lots). The latter parameter represents the portion of the lot conducive to vegetation growth due to soil availability. These area sizes were quantified using the measurement tools available in the municipal GIS system. In cases where the interpretation of aerial imagery within the municipal GIS system for potential area data was ambiguous, verification was conducted via on-lot field inspections. During the process of collecting and analyzing the data, four lots (one per typology C1, D1, E1, and E3) were excluded from the study due to the complete absence of potential space for vegetation growth. Using the three spatial parameters, we also quantified the relative open area per total lot area, and potential area per total lot area.

The assessment of non-maintenance (abandonment) levels for each lot was conducted utilizing five previously-used indicators (Goddard et al., 2017; Loram et al., 2008; Shwartz et al., 2008, 2013). These indicators included (1) the presence of an irrigation system; (2) signs of mowing, pruning, and/or weeding activities; (3) the presence of horticultural plants; (4) the presence of a lawn; and (5) the absence of bare soil. Each indicator was assigned a score ranging from 0 to 1, in 0.5 increments. These scores were summed per lot so that each evaluated yard achieved a total score between 0-5, with a higher score indicating a higher level of non-maintenance.

The proximity of each lot to one of the five largest urban nature areas in the city (those exceeding 30 dunams), which also show the highest species richness (80 species and above) was evaluated. This measurement involved generating circles in AutoCAD software, originating from the center of the sampled lot, and extending to the boundary of the nearest urban nature area.

### Vegetation Sampling

Vegetation sampling within each lot was conducted within the potential open areas, delineated as potential for vegetative growth. Given the variance in conditions across potential areas within a lot (e.g., front versus back, exposure to sunlight and wind), sampling within each lot targeted the section visually assessed as most vegetatively prolific. Within the chosen zone, a 10m-long transect was established for sampling. This transect was aligned along the longest dimension of the selected area. For lots exceeding 2500 m^2^, an additional transect was instituted for every m^2^ of lot area (five lots out of the 56 chosen). The linear transect method was deemed more effective than area sampling (quadrat) due to its enhanced capability to accurately represent the spatial variability present within yard spaces.

Sampling occurred during February-March 2022, a period marked by peak flowering, which facilitated plant identification. Only natural vegetation was surveyed by measuring all individual plants per species within a 25 cm width on each side of the transect. Utilizing this information, both species richness and diversity within each lot were calculated. Species diversity was quantified using the Shannon Index (Shannon & Weiner, 1949):

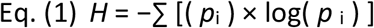

where **pi** is the proportion of individuals belonging to the ith species of the total number of individuals. Values of the Shannon Index can range from 0 to infinity.

To account for potential confounding effects on estimated species richness and diversity due to the multiple transects in lots exceeding 2500 m^2^, we also estimated species richness and diversity using single transects in these lots (the first selected most vegetatively prolific ones). The sampled plant species were also classified into two groups according to their degree of synanthropy: those primarily associated with natural habitats (non-synanthropic) and those primarily associated with human-modified habitats (synanthropic)(Danin, n.d.)

## Statistical analyses

Statistical analyses were performed using IBM SPSS Statistics 29. Variations among the typologies regarding lot sizes, dimensions of open areas, potential area sizes, levels of non-maintenance, species richness, and species diversity were analyzed using Analysis of Variance (ANOVA), with typologies as the fixed factors

The effect of potential area size, non-maintenance levels, and distance from urban nature sites on native plant species richness and diversity were examined using a stepwise multiple regression, with a model selection approach using comparisons of Akaike’s information criterion values (AICc) to decide which variables to enter and remain in the final model (Burnham and Anderson, 2002). Species richness and potential area size were log-transformed prior to analyses to meet parametric assumptions of normality (species richness was log+1 transformed).

## Results

### Lot characterization

The size of the open area per lot varied among typologies, ranging from an average of 248.8 m^2^ in the 150-500 m^2^ box typology to 3544 m^2^ in the tower typology (Figure 3A, Table 1). Similarly, average potential area size varied from 87-3139 m^2^, but was highest in the block typology for lots larger than 1500 m^2^ (Figure 3B, Table 1). The average proportion of open area size across all six typologies was consistently high, accounting for more than half of the lot area in each typology (53-76%) (Figure 3C, Table 1).

**Table 1:** ANOVA results for the effect of lot typology on lot and plant community characteristics.

| Open area size<br>(m<sup>2</sup>) |  | Potential area<br>size (m<sup>2</sup>) |  | open area<br>per lot |  | potential area<br>per lot |  | Non maintenance<br>levels |  |
| --- | --- | --- | --- | --- | --- | --- | --- | --- | --- |
| F | P | F | P | F | P | F | P | F | P |
| 9.327 | < 0.001 | 7.928 | < 0.001 | 9.389 | < 0.001 | 15.291 | < 0.001 | 10.296 | < 0.001 |
| Species diversity |  | Species richness |  | Non-synanthrop<br>species richness |  | Synanthrop<br>Species richness |  |  |  |
| F | P | F | P | F | P | F | P |  |  |
| 2.623 | < 0.035 | 3.927 | 0.004 | 5.478 | < 0.001 | 1.918 | 0.109 |  |  |

characteristics.

**Figure 3:**
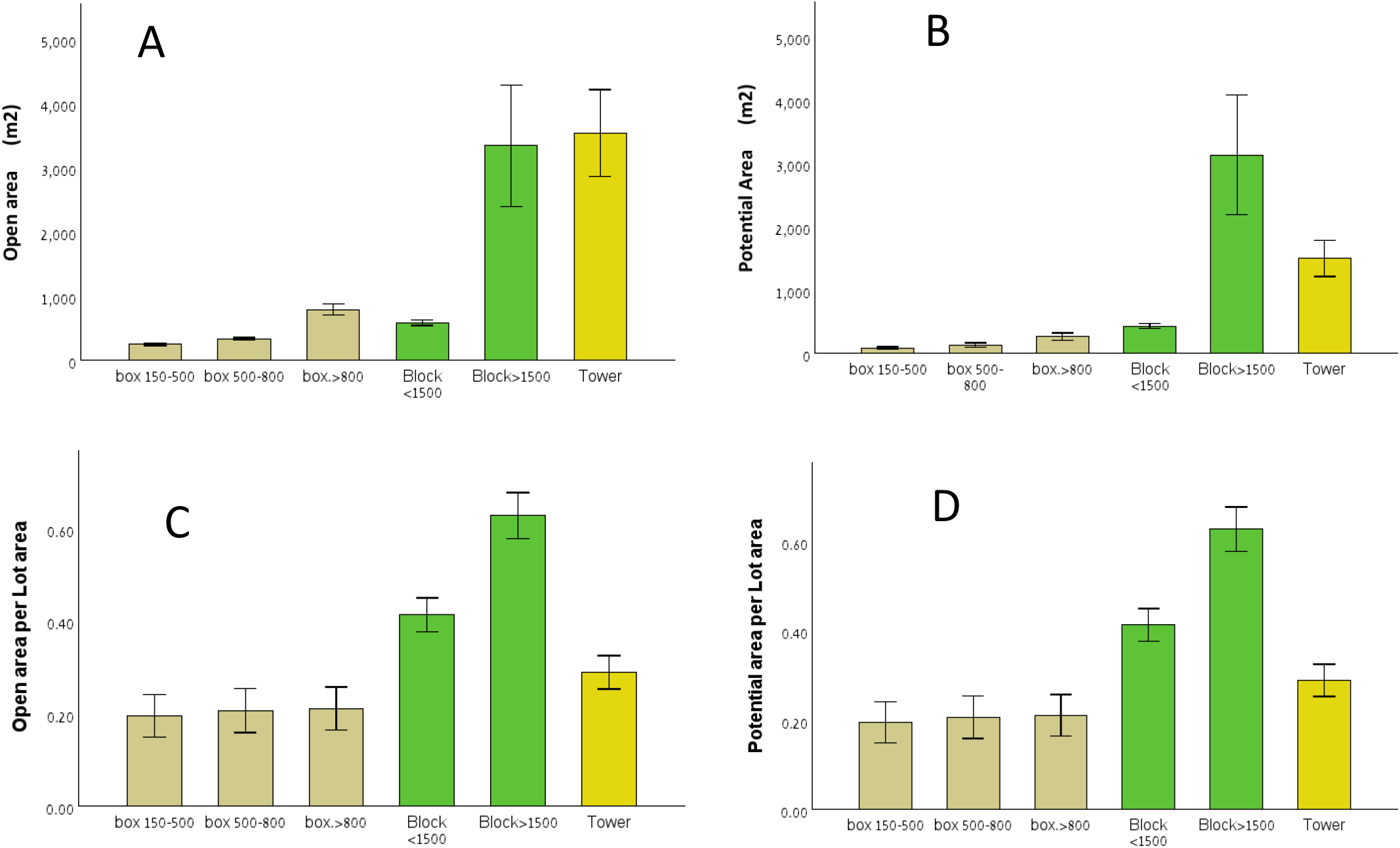
Typology characteristics (means ± SE) according to (A) open area size, (B) potential area size, (C) open area per lot area, and (D) potential area per lot area.

The typologies differed in the proportion of the potential area size relative to the total lot area, ranging from 20% to 62% (Figure 5, Table 1). Specifically, box buildings and towers exhibited the smallest proportion of potential area (20-28%), while block buildings had the highest proportion with potential areas constituting 41-62% of the lot area (Figure 3D, table 1)

**Figure 4:**
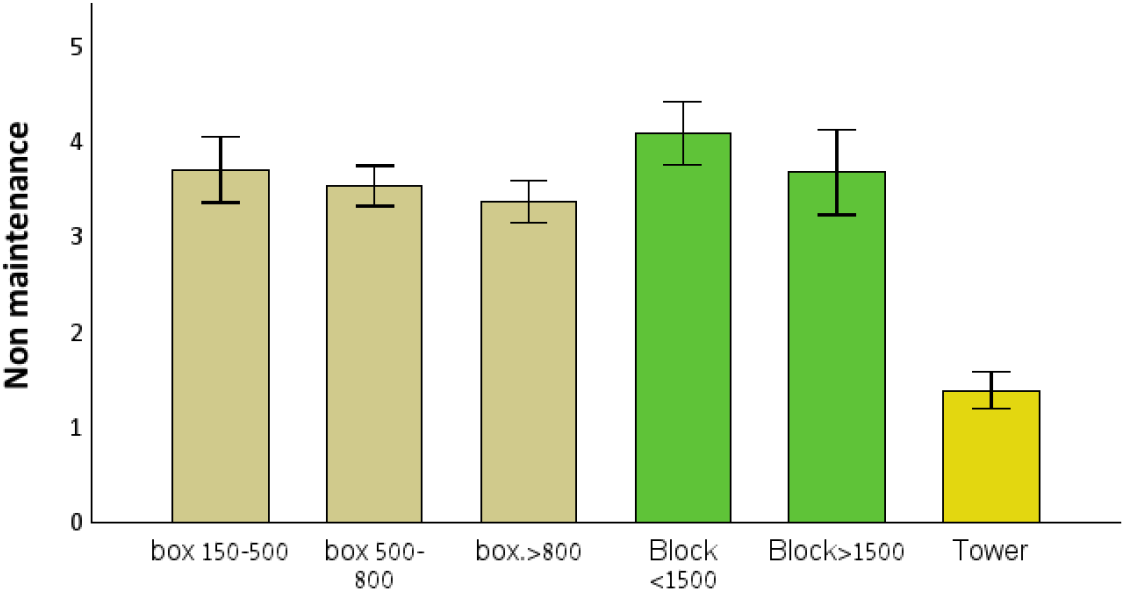
Typology characteristics according to non-maintenance levels (mean ± SE).

**Figure 5:**
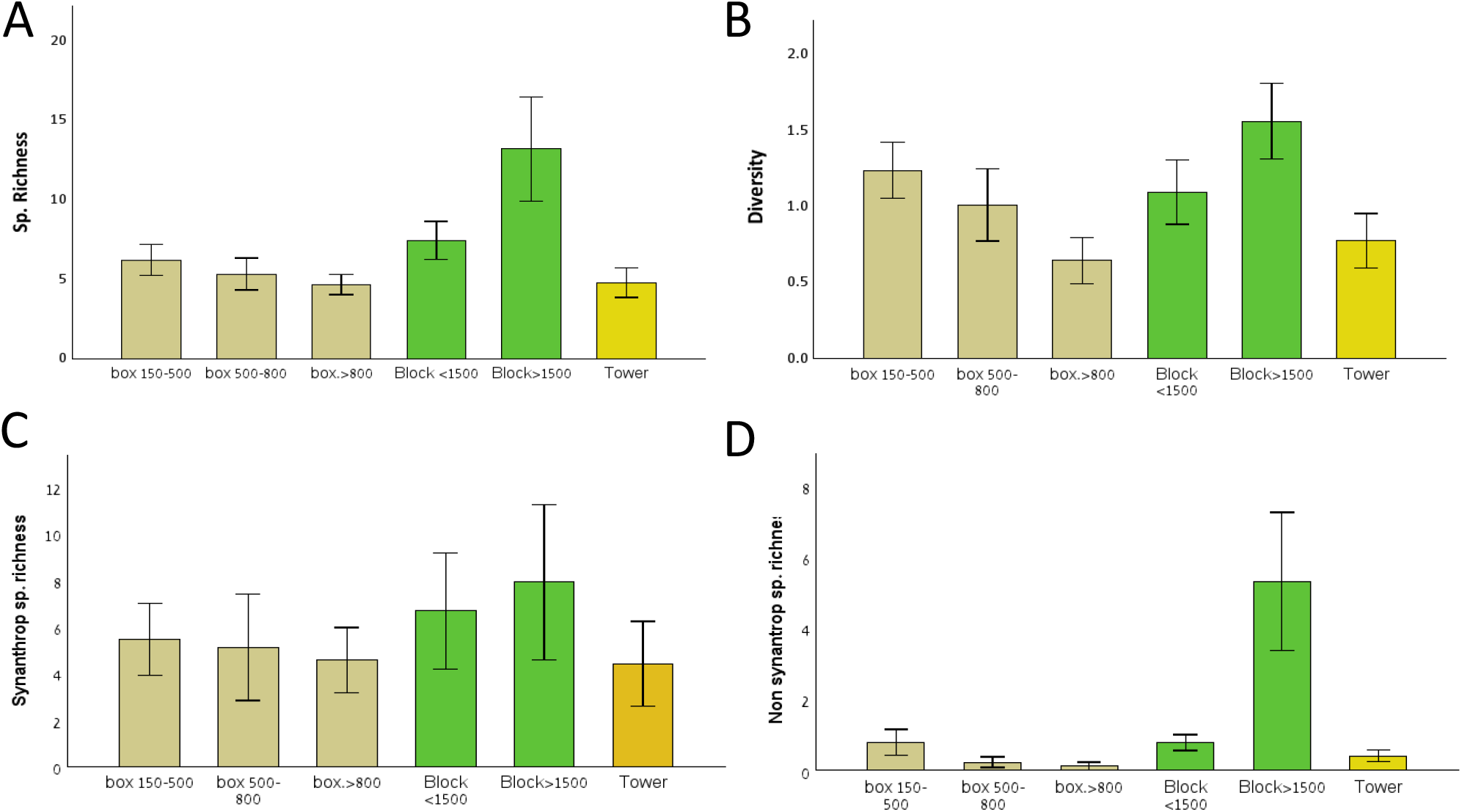
Effects of typology on plant community characteristics (means ± SE), including (A) species richness, (B) species diversity measured as the Shannon Index, (C) synanthropic species richness, and (D) non-synanthropic species richness.

Typologies also differed in their level of non-maintenance (Figure 4, Table 1). The tower typology was distinguished with a low 1.4 non-maintenance level, whereas the block typology on lots smaller than 1500 m^2^ exhibited a high 4.11 level (Figure 4).

### Plant species richness and diversity

A total of 74 wild plant species were identified in the lots. Among these, four species are classified as invasive: *Oxalis pes-caprae, Lantana camara, Asparagus horridus*, and *Conyza bonariensis*. Additionally, nine species were identified as weeds, *including Euphorbia peplus, Euphorbia graminea, Oxalis corniculata, Cardamine occulta, Stellaria pallida, Sagina apetala, Sonchus oleraceus, Senecio vulgaris*, and *Chenopodium chenopodioides*. The documented species encompass mostly native species and a few naturalized aliens. 29 species (39% of the total species identified) were categorized as obligate natural and mostly natural.

The typologies varied in species richness and diversity (Figure 5, Table 1). Specifically lots with a block typology, particularly for lots greater than 1500 m^2^, demonstrated notably high species richness and diversity (Figures 5A,B). Similar results were found when species richness and diversity were estimated using single transects in lots exceeding 2500 m^2^ (species richness: <u>F = 3.54, p = 0.008;</u> species diversity: <u>F = 56.805, p <0.01).</u>

### The effect of potential area and non-maintenance on species richness and diversity

Across typologies, species richness and diversity increased with potential area size (Figure 6A,B, Table 2), and with non-maintenance levels (Figure 6C,D, Table 2). However, distance to the nearest urban nature sites had no effect on species richness and diversity and was dropped from the model (Table 2).

**Table 2:** Results of a stepwise multiple regression for the effect of potential area size, levels of non-maintenance and distance to urban nature sites, on plant species richness and diversity (*P < 0.05, ***P < 0.001).

|  | AIC<sub>c</sub> | R<sup>2</sup> | Intercept | Potential area size | Non maintenance | Distance urban to nature site |
| --- | --- | --- | --- | --- | --- | --- |
| **Species richness** | -2.64 | 0.406 | -0.0677 | 0.1857*** | 0.1251*** | Dropped |
| **Species diversity** | 106.80 | 0.248 | -0.5363 | 0.3179* | 0.2386*** | Dropped |

**Table 3:**
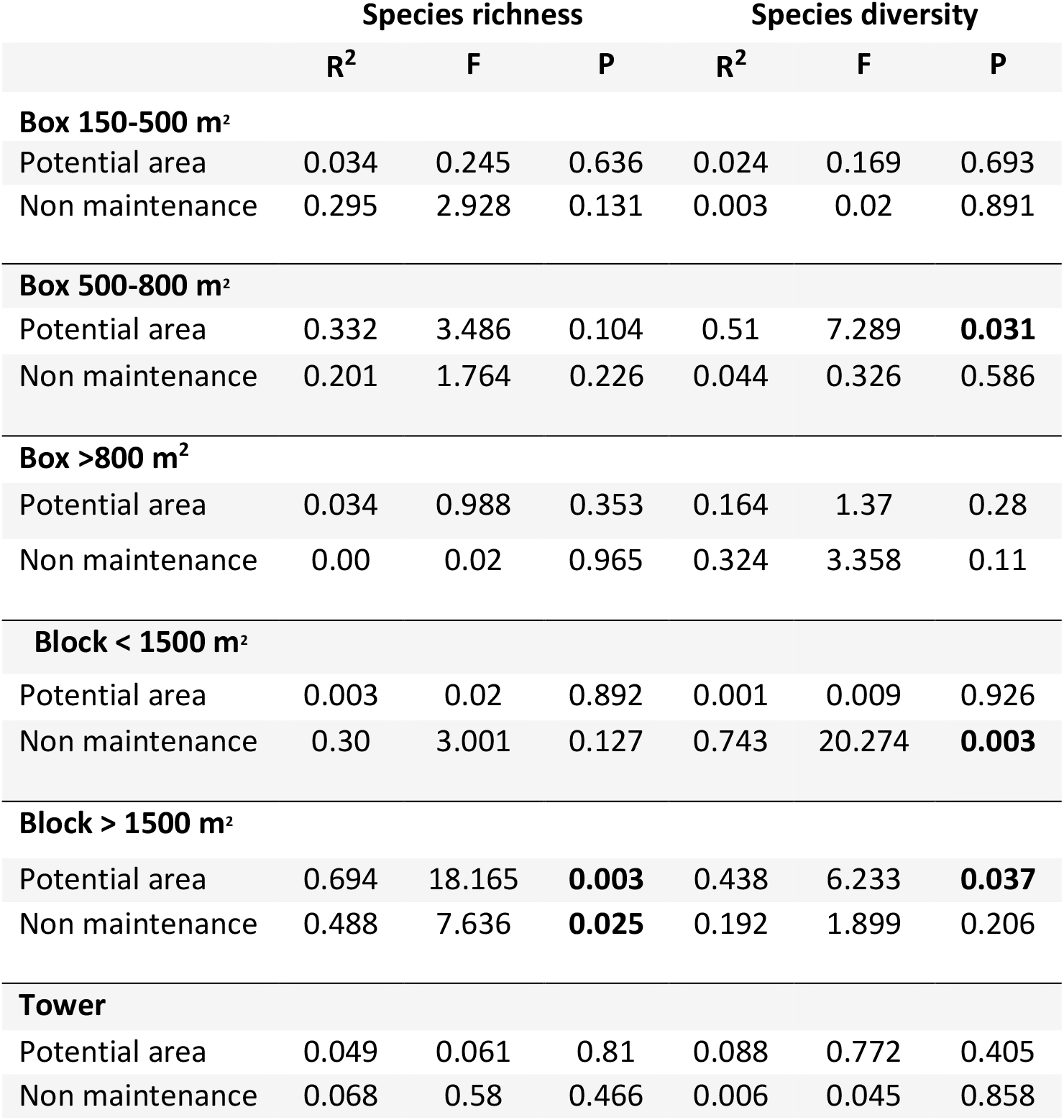
Linear regression results for the effect of potential area and non-maintenance levels on plant species richness and diversity within typology. Significant results are indicated in bold.

|  | Species richness |  |  | Species diversity |  |  |
| --- | --- | --- | --- | --- | --- | --- |
|  | R<sup>2</sup> | F | P | R<sup>2</sup> | F | P |
| **Box 150-500 m<sup>2</sup>** |  |  |  |  |  |  |
| Potential area | 0.034 | 0.245 | 0.636 | 0.024 | 0.169 | 0.693 |
| Non maintenance | 0.295 | 2.928 | 0.131 | 0.003 | 0.02 | 0.891 |
| **Box 500-800 m<sup>2</sup>** |  |  |  |  |  |  |
| Potential area | 0.332 | 3.486 | 0.104 | 0.51 | 7.289 | **0.031** |
| Non maintenance | 0.201 | 1.764 | 0.226 | 0.044 | 0.326 | 0.586 |
| **Box >800 m<sup>2</sup>** |  |  |  |  |  |  |
| Potential area | 0.034 | 0.988 | 0.353 | 0.164 | 1.37 | 0.28 |
| Non maintenance | 0.00 | 0.02 | 0.965 | 0.324 | 3.358 | 0.11 |
| **Block < 1500 m<sup>2</sup>** |  |  |  |  |  |  |
| Potential area | 0.003 | 0.02 | 0.892 | 0.001 | 0.009 | 0.926 |
| Non maintenance | 0.30 | 3.001 | 0.127 | 0.743 | 20.274 | **0.003** |
| **Block > 1500 m<sup>2</sup>** |  |  |  |  |  |  |
| Potential area | 0.694 | 18.165 | **0.003** | 0.438 | 6.233 | **0.037** |
| Non maintenance | 0.488 | 7.636 | **0.025** | 0.192 | 1.899 | 0.206 |
| **Tower** |  |  |  |  |  |  |
| Potential area | 0.049 | 0.061 | 0.81 | 0.088 | 0.772 | 0.405 |
| Non maintenance | 0.068 | 0.58 | 0.466 | 0.006 | 0.045 | 0.858 |

**Figure 6:**
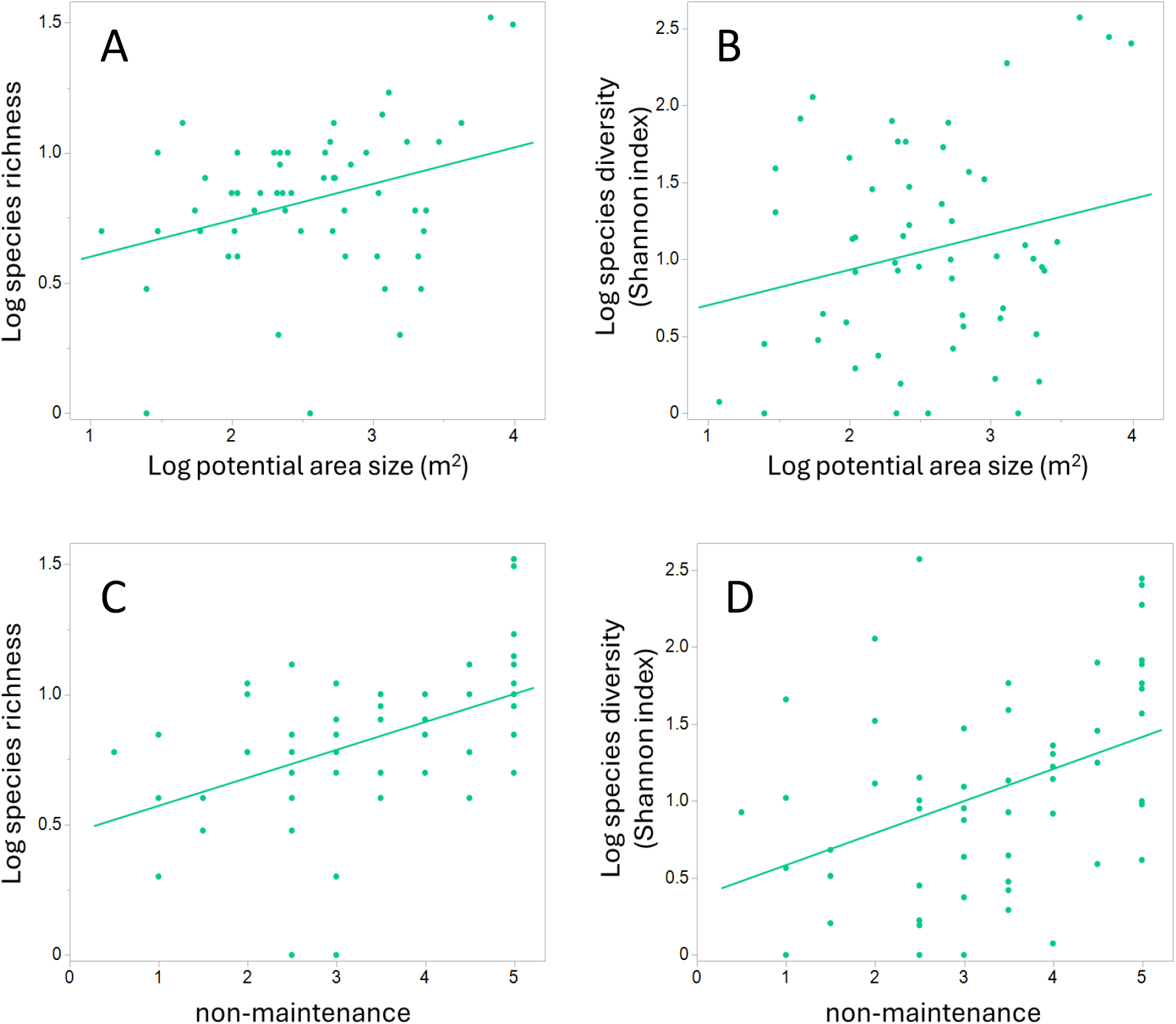
Effects of potential area size and non-maintenance level on (A,C) species richness and (B,D) species diversity.

Within typologies, species richness and diversity were positively affected by the potential area size only in the Block typology on lots bigger than 1500 m^2^. Moreover, all species classified as non-synanthropic were sampled in this typology, making 85% of the species found in it (29 out of the 34 species). As for the effect of potential lot area, species richness within typology increased with the level of non-maintenance only within the Block structures on lots bigger than 1500 m^2^.

## Discussion

In alignment with findings from various urban environments (Goddard et al., 2017; Loram et al., 2008; Mathieu et al., 2007), residential yards in Givatayim constitute a significant portion of the urban landscape, approximately 32%, encompassing around 1,000 dunams. This area significantly exceeds the city’s public open spaces, which total about 220 dunams. Despite their substantial coverage, these green spaces have yet to be fully recognized for their potential to enhance urban natural habitats (Lerman et al., 2023).

This study, based on data collected during the spring of 2022, documented 74 wild plant species across 56 residential yards, representing about a quarter of the total plant species identified in the urban nature survey of the city in 2016. This substantial finding underscores the potential role of residential yards in contributing to the city’s biodiversity. Unlike previous studies focusing on residential private yards, which typically examined the impact of lot characteristics and homeowner preferences (Lerman et al., 2023; Ossola et al., 2019b; Smallwood & Wood, 2023b; Wheeler et al., 2022), this study explores shared yards under collective maintenance, which can significantly affect space management. This study is also distinguished from others by exclusively cataloging wild plant species that have naturally colonized these areas, rather than those deliberately planted by the residents. In addition, in this study we characterized residential yards using a typology system that provided a systematic sampling of different lot sizes and maintenance levels. Using this approach, we found that both lot size and non-maintenance levels contribute positively to the richness and diversity of wild species harbored in the urban environment.

The research highlighted two significant typologies: towers and block buildings situated on lots larger and smaller than 1500 m^2^. Among these, block buildings on larger lots exhibited the highest species richness, including species categorized as non synatropic (i.e. natural with low affinity for humans), which were not observed in other yard typologies. Notably, this specific typology of block buildings on lots larger than 1500 m^2^ was the sole category demonstrating a direct correlation between species richness and potential area size, indicating that these environments offer viable habitats for plant species native to natural settings.

Chronological examination of urban development reveals that, despite towers being the most recent architectural form in terms of construction year, they and the block buildings on lots larger than 1500 m^2^ exhibit similar lot and open area sizes but differ markedly in the size of the potential area for vegetation growth. Specifically, while approximately 70% of the total lot area in both typologies is classified as open space, only 28.9% of this is allocated for gardening in Towers, in contrast to 62.8% in Block buildings. This distinction underscores that, from a planning perspective, the critical metric is not merely the lot size or the open area’s extent, but rather the potential area designated for growth.

The study further identified a correlation between the level of non-maintenance and species richness within the examined lots. Rudolph et al. (2017) highlighted variance in species richness between well-maintained and neglected green spaces. However, they argued that both types of green spaces generally exhibit significantly lower biodiversity compared to semi-natural areas on the urban periphery, and that factors such as soil quality (including acidity, phosphorus levels, etc.) are posited to have a substantial impact on these lower biodiversity outcomes (Rudolph et al., 2017).

A comparison of non-maintenance levels across different typologies revealed a notable disparity with towers, suggesting a shift towards enhanced maintenance practices in such settings (Azian et al., 2020). This inference is supported by the observation that high maintenance costs likely contribute to the limited allocation of softscapes in tower typologies, comprising only 29% of the total lot area. In the block buildings, on the other hand, large areas of soft scapes lead to a characterization of non-maintenance. A thorough exploration of the relationship between non-maintenance practices and architectural typologies necessitates further research beyond the scope of the present study.

The analysis underscores a distinct effect of the potential area size on species richness within the examined samples. This observation aligns with current discussions on urbanization and the focused conservation efforts on large natural expanses, as reported by Deane & He (2018) and Riva & Fahrig (2023). Echoing the sentiments of recent literature, this research emphasizes the critical role of small urban green patches in supporting accessible natural habitats, ecosystem services, and biodiversity (Berthon et al., 2021; Larson et al., 2020b; Lerman et al., 2023; Shwartz et al., 2013). The results reinforce the notion that spatial size directly correlates with both species richness and diversity, even in relatively modest areas ranging from a few hundred to a few thousand square meters. This finding is consistent with a study in Swedish forests (Dynesius & Zinko, 2006), which highlighted a positive relationship between area size and species richness in small natural habitats. However. this raises questions about the minimum size requirement for effective open spaces within urban settings compared to those in natural, undeveloped landscapes.

This insight underscores the critical importance of maintaining diverse-sized green spaces, including small sized ones, in a world of increasing urbanization and population density. Additionally, it highlights a relationship between low-maintenance and enhanced species richness, advocating for a reevaluation of yard maintenance practices to support (Goddard et al., 2017; Lerman et al., 2023; Shwartz et al., 2013). Vega & Kuffer (2021), who examined the influence of the species richness of green areas on each other, determined a distance of 200 m as the maximum range within which areas significantly influence each other. This value is based on the results of Muratet et al. (2007).

Despite these findings, the study does not establish a direct correlation between the proximity of yards to large, diverse urban natural areas and an increase in species richness or diversity in those yards. Research indicates that the characteristics of the lot and its immediate environment have a much greater impact on species richness and diversity within it than the effect of open spaces located hundreds of meters away or more (Turini & Knop 2015; Belaire et al. 2014).

In contrast to larger green spaces, this study found no correlation between species diversity and the size of very small private yards (tens of meters), while a significant relationship emerged in spaces larger than 500 m^2^. This discrepancy prompts a reevaluation of the minimal open space required for effective biodiversity support within urban environments versus natural, undeveloped areas.

## Conclusions

This work contributes to the growing body of research identifying residential yards as vital components in sustaining and enhancing urban biodiversity. It elucidates the connections between maintenance practices, potential area size, and biodiversity, offering a novel approach to mapping urban yards for biodiversity studies and focusing exclusively on wild plant species. The study also acknowledges the critical size threshold for lots supporting diverse species and the importance of soft cover in fostering biodiversity and richness. The relationship between potential area size and species richness underscores the ecological impact of urban green spaces. Amidst urban expansion, current conservation efforts predominantly focus on large natural reserves. However, this research, echoing recent work, advocates for the preservation of smaller urban green patches to support accessible nature, ecosystem services, and biodiversity. Empirical evidence confirms that even compact areas exhibit a positive correlation with biodiversity. This highlights the importance of integrating small-scale green spaces into urban planning and conservation strategies.

In the examination of the relationship between species richness/biodiversity and the extent of non-maintenance, significance was identified exclusively within block-associated lots. These observations highlight a prominent feature of both categories: a substantial proportion of potential area relative to total area coupled with significant non-maintenance. Towers contrast this pattern by exhibiting a high maintenance approach.

The study also reinforces the notion that low-maintenance areas contribute to species richness, challenging traditional yard maintenance perspectives. However, it did not establish a direct link between yard proximity to large urban natural areas and increased species diversity, suggesting that even small, less maintained areas play a crucial role in urban biodiversity.

Aligning with recent research, this study highlights residential yards as vital for enhancing urban biodiversity, revealing significant links between minimal maintenance and species richness, as well as the relationship between the size of potential areas and plant diversity. Distinguishing itself by focusing on shared yards and developing a method for city-wide yard mapping for wild plant sampling, it underscores the importance of certain lot sizes for sustaining diverse plant life. This approach not only broadens our understanding of urban green spaces but also emphasizes the ecological value of less maintained areas.

## Bibliography

Aronson, M. F., Lepczyk, C. A., Evans, K. L., Goddard, M. A., Lerman, S. B., MacIvor, J. S., Nilon, C. H., & Vargo, T. (2017). Biodiversity in the city: key challenges for urban green space management [Article]. Frontiers in Ecology and the Environment, 15(4), 189–196. 10.1002/fee.1480

Azian, F. U. M., Yusof, N., & Kamal, E. M. (2020). Problems in high rise residential building: From management perspective [Article]. IOP Conference Series: Earth and Environmental Science, 452(1), 12087. 10.1088/1755-1315/452/1/012087

Baldock, K. C. R., Goddard, M. A., Hicks, D. M., Kunin, W. E., Mitschunas, N., Morse, H., Osgathorpe, L. M., Potts, S. G., Robertson, K. M., Scott, A. V, Staniczenko, P. P. A., Stone, G. N., Vaughan, I. P., & Memmott, J. (2019). A systems approach reveals urban pollinator hotspots and conservation opportunities. Nature Ecology and Evolution, 3(3), 363–373. 10.1038/s41559-018-0769-y

Beninde, J., Veith, M., & Hochkirch, A. (2015). Biodiversity in cities needs space: a meta-analysis of factors determining intra-urban biodiversity variation [Article]. Ecology Letters, 18(6), 581–592. 10.1111/ele.12427

Berghauser Pont, M., Stavroulaki, G., Bobkova, E., Gil, J., Marcus, L., Olsson, J., Sun, K., Serra, M., Hausleitner, B., Dhanani, A., & Legeby, A. (2019). The spatial distribution and frequency of street, plot and building types across five European cities [Article]. Environment and Planning. B, Urban Analytics and City Science, 46(7), 1226–1242. 10.1177/2399808319857450

Berthon, K., Thomas, F., & Bekessy, S. (2021). The role of ‘nativeness’ in urban greening to support animal biodiversity [Article]. Landscape and Urban Planning, 205, 103959. 10.1016/j.landurbplan.2020.103959

Beumer, C., & Martens, P. (2015). Biodiversity in my (back)yard: towards a framework for citizen engagement in exploring biodiversity and ecosystem services in residential gardens. Sustainability Science, 10(1), 87–100. 10.1007/s11625-014-0270-8

Blanchette, A., Trammell, T. L. E., Pataki, D. E., Endter-Wada, J., & Avolio, M. L. (2021). Plant biodiversity in residential yards is influenced by people’s preferences for variety but limited by their income. Landscape and Urban Planning, 214. 10.1016/j.landurbplan.2021.104149

Botzat, A., Fischer, L. K., & Kowarik, I. (2016). Unexploited opportunities in understanding liveable and biodiverse cities. A review on urban biodiversity perception and valuation. Global Environmental Change, 39, 220–233. 10.1016/j.gloenvcha.2016.04.008

Burghardt, K. T., Tallamy, D. W., & Gregory Shriver, W. (2009). Impact of Native Plants on Bird and Butterfly Biodiversity in Suburban Landscapes [Article]. Conservation Biology, 23(1), 219–224. 10.1111/j.1523-1739.2008.01076.x

Chamberlain, D. E., Gough, S., Vaughan, H., Vickery, J. A., & Appleton, G. F. (2007). Determinants of bird species richness in public green spaces [Article]. Bird Study, 54(1), 87–97. 10.1080/00063650709461460

Chen, F. (2012). Interpreting urban micromorphology in China: case studies from Suzhou [Article]. Urban Morphology, 16(2), 133–148. 10.51347/jum.v16i2.3985

Coetzee, A., Barnard, P., & Pauw, A. (2018). Urban nectarivorous bird communities in Cape Town, South Africa, are structured by ecological generalisation and resource distribution [Article]. Journal of Avian Biology, 49(6), n/a. 10.1111/jav.01526

Colding, J. (2007). “Ecological land-use complementation” for building resilience in urban ecosystems. Landscape and Urban Planning, 81(1–2), 46–55. 10.1016/j.landurbplan.2006.10.016

Danin, A. and. F.-S. O. (n.d.). Flora of Israel and adjacent areas. Retrieved July 9, 2024, from https://flora.org.il/en/plants/

Deane, D. C., & He, F. (2018). Loss of only the smallest patches will reduce species diversity in most discrete habitat networks. Global Change Biology, 24(12), 5802–5814. 10.1111/gcb.14452

Dynesius, M., & Zinko, U. (2006). Species richness correlations among primary producers in boreal forests [Article]. Diversity & Distributions, 12(6), 703–713. 10.1111/j.1472-4642.2006.00280.x

Fahrig, L., & Storch, D. (2020). Why do several small patches hold more species than few large patches? [Article]. Global Ecology and Biogeography, 29(4), 615–628. 10.1111/geb.13059

Filazzola, A., Shrestha, N., & MacIvor, J. S. (2019). The contribution of constructed green infrastructure to urban biodiversity: A synthesis and meta-analysis. In Journal of Applied Ecology (Vol. 56, Issue 9, pp. 2131–2143). Blackwell Publishing Ltd. 10.1111/1365-2664.13475

Fuller, R. A., Irvine, K. N., Devine-Wright, P., Warren, P. H., & Gaston, K. J. (2007). Psychological benefits of greenspace increase with biodiversity. Biology Letters, 3(4), 390–394. 10.1098/rsbl.2007.0149

Gerner, E. E., & Sargent, R. D. (2022a). Local plant richness predicts bee abundance and diversity in a study of urban residential yards. Basic and Applied Ecology, 58, 64–73. 10.1016/j.baae.2021.11.004

Gerner, E. E., & Sargent, R. D. (2022b). Local plant richness predicts bee abundance and diversity in a study of urban residential yards. Basic and Applied Ecology, 58, 64–73. 10.1016/J.BAAE.2021.11.004

Goddard, M. A., Ikin, K., & Lerman, S. B. (2017). Ecological and social factors determining the diversity of birds in residential yards and gardens. In Ecology and Conservation of Birds in Urban Environments (pp. 371–397). Springer International Publishing. 10.1007/978-3-319-43314-1_18

Grade, A. M., Lerman, S. B., & Warren, P. S. (2021). Perilous choices: landscapes of fear for adult birds reduces nestling condition across an urban gradient. Ecosphere, 12(7). 10.1002/ecs2.3665

Hammill, E., Clements, C. F., & Hodgson, D. (2020). Imperfect detection alters the outcome of management strategies for protected areas [Article]. Ecology Letters, 23(4), 682–691. 10.1111/ele.13475

Hobbs, R. J., Arico, S., Aronson, J., Baron, J. S., Bridgewater, P., Cramer, V. A., Epstein, P. R., Ewel, J. J., Klink, C. A., Lugo, A. E., Norton, D., Ojima, D., Richardson, D. M., Sanderson, E. W., Valladares, F., Vilà, M., Zamora, R., & Zobel, M. (2006). Novel ecosystems: Theoretical and management aspects of the new ecological world order. Global Ecology and Biogeography, 15(1), 1–7. 10.1111/j.1466-822X.2006.00212.x

Hoelzer, G. A., Smith, E., & Pepper, J. W. (2006). On the logical relationship between natural selection and self-organization. Journal of Evolutionary Biology, 19(6), 1785–1794. 10.1111/j.1420-9101.2006.01177.x

Israel Meteorological Service. (n.d.). Retrieved July 9, 2024, from https://ims.gov.il/en

Jimenez, M. F., Pejchar, L., Reed, S. E., & McHale, M. R. (2022). The efficacy of urban habitat enhancement programs for conserving native plants and human-sensitive animals [Article]. Landscape and Urban Planning, 220, 104356. 10.1016/j.landurbplan.2022.104356

Kevin, G. J. (2008). Urban domestic gardens (XII): The richness and composition of the flora in five UK cities. Journal of Vegetation Science, 19, 321–330. 10.3170/2007-8-18373

Kremen, C. (2015). Reframing the land-sparing/land-sharing debate for biodiversity conservation [Article]. Annals of the New York Academy of Sciences, 1355(1), 52–76. 10.1111/nyas.12845

Larsen, L., & Harlan, S. L. (2006). Desert dreamscapes: Residential landscape preference and behavior. Landscape and Urban Planning, 78(1–2), 85–100. 10.1016/j.landurbplan.2005.06.002

Larson, K. L., Andrade, R., Nelson, K. C., Wheeler, M. M., Engebreston, J. M., Hall, S. J., Avolio, M. L., Groffman, P. M., Grove, M., Heffernan, J. B., Hobbie, S. E., Lerman, S. B., Locke, D. H., Neill, C., Chowdhury, R. R., & Trammell, T. L. E. (2020a). Municipal regulation of residential landscapes across US cities: Patterns and implications for landscape sustainability. Journal of Environmental Management, 275. 10.1016/j.jenvman.2020.111132

Larson, K. L., Andrade, R., Nelson, K. C., Wheeler, M. M., Engebreston, J. M., Hall, S. J., Avolio, M. L., Groffman, P. M., Grove, M., Heffernan, J. B., Hobbie, S. E., Lerman, S. B., Locke, D. H., Neill, C., Chowdhury, R. R., & Trammell, T. L. E. (2020b). Municipal regulation of residential landscapes across US cities: Patterns and implications for landscape sustainability. Journal of Environmental Management, 275, 111132. 10.1016/J.JENVMAN.2020.111132

Larson, K. L., Lerman, S. B., Nelson, K. C., Narango, D. L., Wheeler, M. M., Groffman, P. M., Hall, S. J., & Grove, J. M. (2022). Examining the potential to expand wildlife-supporting residential yards and gardens. Landscape and Urban Planning, 222. 10.1016/j.landurbplan.2022.104396

Lerman, S. B., Larson, K. L., Narango, D. L., Goddard, M. A., & Marra, P. P. (2023). Humanity for Habitat: Residential Yards as an Opportunity for Biodiversity Conservation [Article]. Bioscience, 73(9), 671–689. 10.1093/biosci/biad085

Lerman, S. B., & Warren, P. S. (2011). The conservation value of residential yards: Linking birds and people. Ecological Applications, 21(4), 1327–1339. 10.1890/10-0423.1

Lin, B. B., & Fuller, R. A. (2013). FORUM: Sharing or sparing? How should we grow the world’s cities? Journal of Applied Ecology, 50(5), 1161–1168. 10.1111/1365-2664.12118

Loram, A., Warren, P. H., & Gaston, K. J. (2008). Urban domestic gardens (XIV): The characteristics of gardens in five cities. Environmental Management, 42(3), 361–376. 10.1007/s00267-008-9097-3

Mathieu, R., Freeman, C., & Aryal, J. (2007). Mapping private gardens in urban areas using object-oriented techniques and very high-resolution satellite imagery. Landscape and Urban Planning, 81(3), 179–192. 10.1016/J.LANDURBPLAN.2006.11.009

Mcdonald, R. I., Kareiva, P., & Forman, R. T. T. (2008). The implications of current and future urbanization for global protected areas and biodiversity conservation. Biological Conservation, 141(6), 1695–1703. 10.1016/j.biocon.2008.04.025

Mendelson, A. (2016). Urban natural infrastructure survey Givatayim (In Hebrew). https://www.givatayim.muni.il/uploads/n/1602745428.4464.pdf

Murdoch, R., & Al-Habashna, A. (2024). Residential building type classification from street-view imagery with convolutional neural networks [Article]. Signal, Image and Video Processing, 18(2), 1949–1958. 10.1007/s11760-023-02882-8

Niemelä, J. (1999). Ecology and urban planning. Biodiversity and Conservation, 8(1), 119–131. 10.1023/A:1008817325994

Ossola, A., Locke, D., Lin, B., & Minor, E. (2019a). Greening in style: Urban form, architecture and the structure of front and backyard vegetation. Landscape and Urban Planning, 185, 141–157. 10.1016/j.landurbplan.2019.02.014

Ossola, A., Locke, D., Lin, B., & Minor, E. (2019b). Yards increase forest connectivity in urban landscapes. Landscape Ecology, 34(12), 2935–2948. 10.1007/s10980-019-00923-7

Rabinowitz, D. (2008). Private Apartment, Shared Building, Public Space: The Israeli Residential Unit Between Personal Space and Public Domain. In S. Cohen & T. Amir (Eds.), Shapes of living, Architecture, and Society in Israel: Vol. In Hebrew (pp. 145–166). Xargol Books LTD & Am oved publishers.

Ripplinger, J., Collins, S. L., York, A. M., & Franklin, J. (2017). Boom–bust economics and vegetation dynamics in a desert city: How strong is the link? [Article]. Ecosphere (Washington, D.C), 8(5), n/a. 10.1002/ecs2.1826

Riva, F., & Fahrig, L. (2023). Obstruction of biodiversity conservation by minimum patch size criteria [Article]. Conservation Biology, 37(5), e14092–e14092. 10.1111/cobi.14092

Rudolph, M., Velbert, F., Schwenzfeier, S., Kleinebecker, T., & Klaus, V. H. (2017). Patterns and potentials of plant species richness in high-and low-maintenance urban grasslands [Article]. Applied Vegetation Science, 20(1), 18–27. 10.1111/avsc.12267

Shanahan, D. F., Miller, C., Possingham, H. P., & Fuller, R. A. (2011). The influence of patch area and connectivity on avian communities in urban revegetation. Biological Conservation, 144(2), 722–729. 10.1016/j.biocon.2010.10.014

Shannon, C. E., & Weiner, V. (1949). A Mathematical Theory of Communication University Press. Illinois Urban, 1, 101–107.

Shwartz, A., Muratet, A., Simon, L., & Julliard, R. (2013). Local and management variables outweigh landscape effects in enhancing the diversity of different taxa in a big metropolis. Biological Conservation, 157, 285–292. 10.1016/J.BIOCON.2012.09.009

Shwartz, A., Shirley, S., & Kark, S. (2008). How do habitat variability and management regime shape the spatial heterogeneity of birds within a large Mediterranean urban park? [Article]. Landscape and Urban Planning, 84(3), 219–229. 10.1016/j.landurbplan.2007.08.003

Smallwood, N. L., & Wood, E. M. (2023a). The ecological role of native-plant landscaping in residential yards to birds during the nonbreeding period. Ecosphere, 14(1). 10.1002/ecs2.4360

Smallwood, N. L., & Wood, E. M. (2023b). The ecological role of native-plant landscaping in residential yards to birds during the nonbreeding period. Ecosphere, 14(1). 10.1002/ecs2.4360

Soga, M., Yamaura, Y., Koike, S., & Gaston, K. J. (2014). Land sharing vs. land sparing: Does the compact city reconcile urban development and biodiversity conservation? Journal of Applied Ecology, 51(5), 1378–1386. 10.1111/1365-2664.12280

Tooke, T. R., vanderLaan, M., Coops, N., Christen, A., & Kellett, R. (2011). Classification of residential building architectural typologies using LiDAR [Proceeding]. 2011 Joint Urban Remote Sensing Event, 221–224. 10.1109/JURSE.2011.5764760

Valente, J. J., Bennett, R. E., Gómez, C., Bayly, N. J., Rice, R. A., Marra, P. P., Ryder, T. B., & Sillett, T. S. (2022). Land-sparing and land-sharing provide complementary benefits for conserving avian biodiversity in coffee-growing landscapes [Article]. Biological Conservation, 270, 109568. 10.1016/j.biocon.2022.109568

Wheeler, M. M., Larson, K. L., Bergman, D., & Hall, S. J. (2022). Environmental attitudes predict native plant abundance in residential yards. Landscape and Urban Planning, 224. 10.1016/j.landurbplan.2022.104443

Wood, B., Mcmi, D., Mrics, R., Fciob, M., & Wiley-Blackwell,). (n.d.). Building Maintenance.

Zhang, J. (2014). Urban morphological processes in China: a Conzenian approach [Article]. Urban Morphology, 19(1), 35–56. 10.51347/jum.v19i1.4023

